# Polarized α-synuclein trafficking and transcytosis across Brain Endothelial Cells via Rab7-decorated carriers

**DOI:** 10.1101/2021.12.21.473642

**Authors:** Parvez Alam, Mikkel R. Holst, Line Lauritsen, Janni Nielsen, Simone S. E. Nielsen, Poul Henning Jensen, Jonathan R. Brewer, Daniel E. Otzen, Morten S. Nielsen

**Author notes:** **Address and Email for correspondence:** Morten S. Nielsen, Department of Biomedicine, Faculty of Health, Aarhus University, Aarhus C, Denmark or Daniel E. Otzen, iNANO, Aarhus University, Aarhus C, Denmark. Equal contributors (Parvez Alam, Mikkel R. Holst). **Author Contributions:** PA, MRH, DO and MSN conceived the study. PA, MRH, JB, PH, DO, and MSN designed experiments. PA, MRH, LL, SN and JN performed experiments. PA, MRH and LL analyzed data. PA and MRH wrote the manuscript with input from SN, DO and MSN. **Competing Interest Statement:** Authors declare no competing interests.

## Abstract

Parkinson’s disease is mainly caused by aggregation of α-synuclein (α-syn) in the brain. Exchange of α-syn between the brain and peripheral tissues could have important pathophysiological and therapeutic implications, but the trafficking mechanism of α-syn across the blood brain barrier (BBB) remains unclear. In this study, we therefore investigated uptake and transport mechanisms of α-syn monomers and oligomers across an *in vitro* BBB model system. Both α-syn monomers and oligomers were internalized by primary brain endothelial cells, with increased restriction of oligomeric over monomeric transport. To enlighten the trafficking route of monomeric α-syn in brain endothelial cells, we investigated co-localization of α-syn and intracellular markers of vesicular transport. Here, we observed the highest colocalization with clathrin, Rab7 and VPS35, suggesting a clathrin-dependent internalization, preferentially followed by a late endosome retromer-connected trafficking pathway. Furthermore, STED microscopy revealed monomeric α-syn trafficking via Rab7-decorated carriers. Knockdown of Caveolin1, VPS35, and Rab7 using siRNA did not affect monomeric α-syn uptake into endothelial cells. However, it significantly reduced transcytosis of monomeric α-syn in the luminal-abluminal direction, suggesting a polarized regulation of monomeric α-syn vesicular transport. Our findings suggest a direct role for Rab7 in polarized trafficking of monomeric α-syn across BBB endothelium, and the potential of Rab7 directed trafficking to constitute a target pathway for new therapeutic strategies against Parkinson’s disease and related synucleinopathies.

**Significance Statement:** In the submitted manuscript, we describe the use of a state-of-the-art porcine blood-brain barrier model based on primary cells to get information about this important issue. We identify several hitherto undescribed cellular pathways to mediate polarized transport of alpha-synuclein. One of these paths we find regulated by Rab7 and can be inhibited by targeting several intracellular proteins such as VPS35, Caveolin1 and Rab7. New knowledge describing brain endothelial intracellular transport systems are highly warranted for identifying new target to alleviate Parkinson’s disease. We believe that our findings could be the seed to establish new therapeutic strategies against Parkinson’s disease and related synucleinopathies.

## Introduction

Parkinson’s disease (PD) is the second most common neurodegenerative disease, characterized by loss of dopaminergic neurons and presence of intracellular inclusions called Lewy bodies throughout the brain (1, 2). The main protein component of Lewy bodies is the 140-residue protein α-synuclein (α-syn), which to a large extend is expressed in presynaptic neurons and to a less extent in other parts of the body (3). While the exact physiological role of α-syn remains unclear, it is reported to be implicated in synaptic formation, regulation of presynaptic vesicle pool, release of neurotransmitter, exocytosis, nerve cell adhesion, synaptic function, and plasticity (4).

Under physiological conditions, α-syn has no persistent structure in the monomeric state. However, as intimated by its presence in Lewy bodies, α-syn has an intrinsic tendency to form aggregates of higher molecular weights, including oligomeric species and fibrils. This tendency is exacerbated by physiological challenges such as pH variation, oxidative stress, mutations and over-expression of the SNCA gene (5). Numerous α-syn oligomeric species have been reported to vary in morphology, structure and molecular weight (6), and can be categorized into on- and off-pathway oligomers. On-pathway oligomers are directly incorporated into fibrillar assemblies, while off-pathway oligomers do not form fibrils (7, 8) and are often sufficiently stable for purification by chromatography (9). α-syn oligomers exert many pathogenic effects including cytoskeletal alterations, membrane permeabilization, increased Ca^2+^ influx, increased reactive oxygen species, decreased neuronal excitability, decreased synaptic firing and impaired protein degradation and turnover (10-13). α-syn is considered an intracellular protein, but it has also been found in blood and cerebrospinal fluid (CSF) (14, 15). Decreased levels of total α-syn in CSF have been reported for PD patients compared to healthy controls (16). Remarkably, high levels of oligomeric α-syn have been reported in CSF of PD patients compared to healthy controls (17). Such accumulation of oligomeric α-syn may be accounted for by defective transport mechanisms across the blood brain barrier (BBB), but it remains unclear with the transport mechanism of α-syn not yet being fully understood.

The BBB form a highly selective, semi-permeable interface between the central nervous system (CNS) and systemic circulation. It allows controlled exchange of essential molecules such as sugars, amino acids, and lipids, required for proper synaptic and neuronal functioning and brain homeostasis (18). The BBB also acts as gatekeeper to protect brain cells from external toxic factors in the systemic circulation, a function that is maintained by specialized efflux transporters (19). This physical and functional regulation is maintained by the intricate interaction between the brain endothelial cells (BECs), astrocytes and pericytes of the BBB, together referred to as the neurovascular unit. (20). BECs have specialized tight- and adherens junctions, restricting the paracellular transport and thereby transfer of substances between the CNS and blood (21). Disruption of tight junctions may lead to BBB breakdown, which has been reported in Alzheimer’s as well as other neurodegenerative diseases (20). Such breakdown can disrupt the controlled transport between blood and brain parenchyma. Several proteins involved in neurodegenerative disorders are believed to be transported across the BBB by controlled transport paths. Amyloid β peptide, the main component of amyloid plaques in Alzheimer’s disease patients, crosses the BBB in both directions (blood to brain and *vice versa*) (22). Likewise, *in vivo* studies have shown that α-syn crosses the BBB in both directions (23). A more recent study has demonstrated the transport of α-syn-containing extracellular vesicles (derived from erythrocytes) across BBB in a process that involves an adsorptive mechanism of transcytosis (24). It has been reported earlier that α-syn is released by neurons into interstitial fluid that merges with CSF (25). The released α-syn is readily taken up by astrocytes, inducing formation of pathological inclusions and degenerative changes (26). α-syn aggregated species have also been found in astrocytes from post mortem PD patient brains (27).

Though α-syn is expressed by blood cells and is found in the brain, little is known about the cellular mechanisms by which α-syn is transported in and out of the brain. A better understanding of the mechanism of transport of monomeric and oligomeric α-syn species through BECs may provide important insights into the normal physiological and pathological regulation of α-syn. To address this, we here report a study of the transport and uptake of α-syn monomers and oligomers across the BBB using an *in vitro* model generated from BECs of porcine origin (pBECs) and astrocytes of rat origin. We determine the preferred polarized trafficking route of monomeric α-syn across the endothelial cells in both directions (from luminal to abluminal and from abluminal to luminal – mimicking blood to brain and brain to blood transport, respectively) (28). We find that both monomeric and oligomeric α-syn is regulated by polarized transport systems. The polarized trafficking route of monomeric α-syn in BECs was dissected by performing co-staining of α-syn with various endo-lysosomal markers. Knock-down of Caveolin1, VPS35 and Rab7 using siRNA confirmed their role in trafficking of monomeric α-syn. Colocalization of α-syn with Rab7 was further pinpointed by STED microscopy that highlights the direct role of Rab7 in polarized trafficking of α-syn across the BBB.

## Results

### Characterization of BBB model

For this study we used a previously described BBB model based on pBECs established in non-contact co-culture (NCC) with rat astrocytes (see setup in **Fig. 1a**) (28). The integrity of the endothelial cell monolayer grown in NCC was assessed by measuring TEER. The obtained average TEER values at the day of the experiments was 1233+36 Ω*cm^2^ (**Fig. 1b**) which indicates a barrier tight enough to stop molecules as small as 521-Da Luciferase Yellow (α-syn is 14.46 kDa(28)). Analysis of staining for tight junction marker protein showed the expected presence of the tight junction marker proteins claudin 5 and ZO-1 (**Fig. 1c-d**). Overall, these results confirmed the quality and tightness of the model and thereby its suitability for studying transcellular transport of α-syn across pBECs.

**Figure 1:**
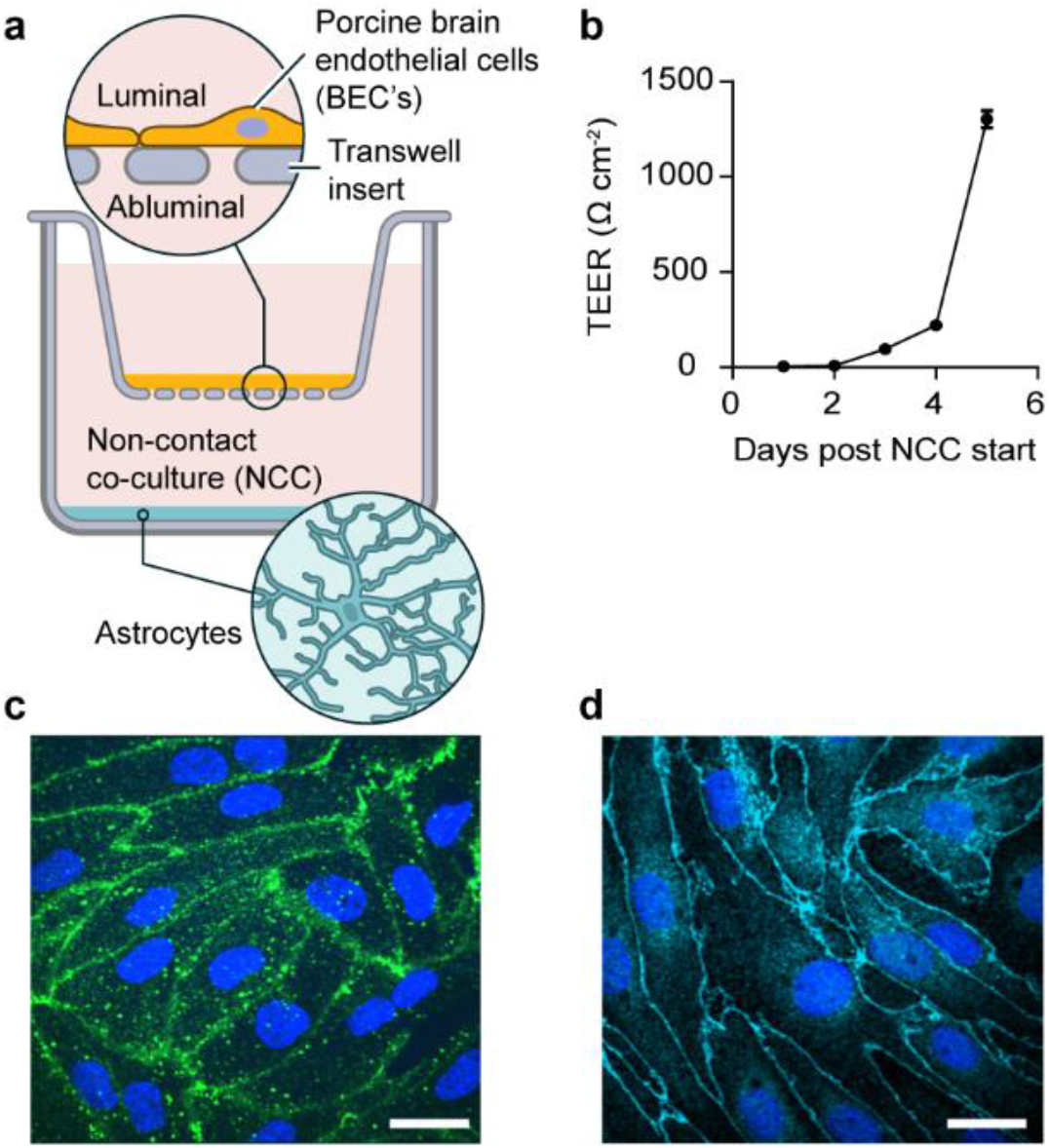
The experimental BBB model. Schematic for the *in vitro* BBB model of pBECs (porcine brain endothelial cells) in Non-Contact Co-culture (NCC) with astrocytes (a). Validation of the model with representative TEER (trans endothelial cells resistance) values post NCC days (b). Representative micrographs of immunostainings for tight junction marker proteins showing (c) Claudin 5 (c) and ZO-1 (d). Scale bars show 15 µm.

### Uptake and transport of α-syn monomers and oligomers across BBB model

To investigate α-syn uptake, we added α-syn monomers from the luminal or abluminal side of Transwell seeded pBECs respectively, and fixed the cells after 10 min or 2 h chase. Confocal imaging revealed binding and uptake of α-syn within 10 min, which was maintained after 2 h. Each setup resulted in clear uptake of α-syn, but the amount endocytosed and the subsequent localization appeared more homogenous after 2 h of chase (**Fig. 2a-d**). We also measured the amount of α-syn crossing the endothelial cell barrier (see schematics in **Fig. 2e**). Using an ELISA setup, we found no measurable transport of α-syn from one side to the other within 10 min, irrespective of the treatment side (**Fig. 2f-g**, acceptor side). However, after 2 h chase, a significant amount of α-syn had crossed the barrier from the abluminal to the luminal compartment (68.60 ± 3.86) % (**Fig. 2f**) and slightly smaller amounts in the other direction (53.92 ± 5.88%) (**Fig. 2g** and **Fig. S1**).

**Figure 2:**
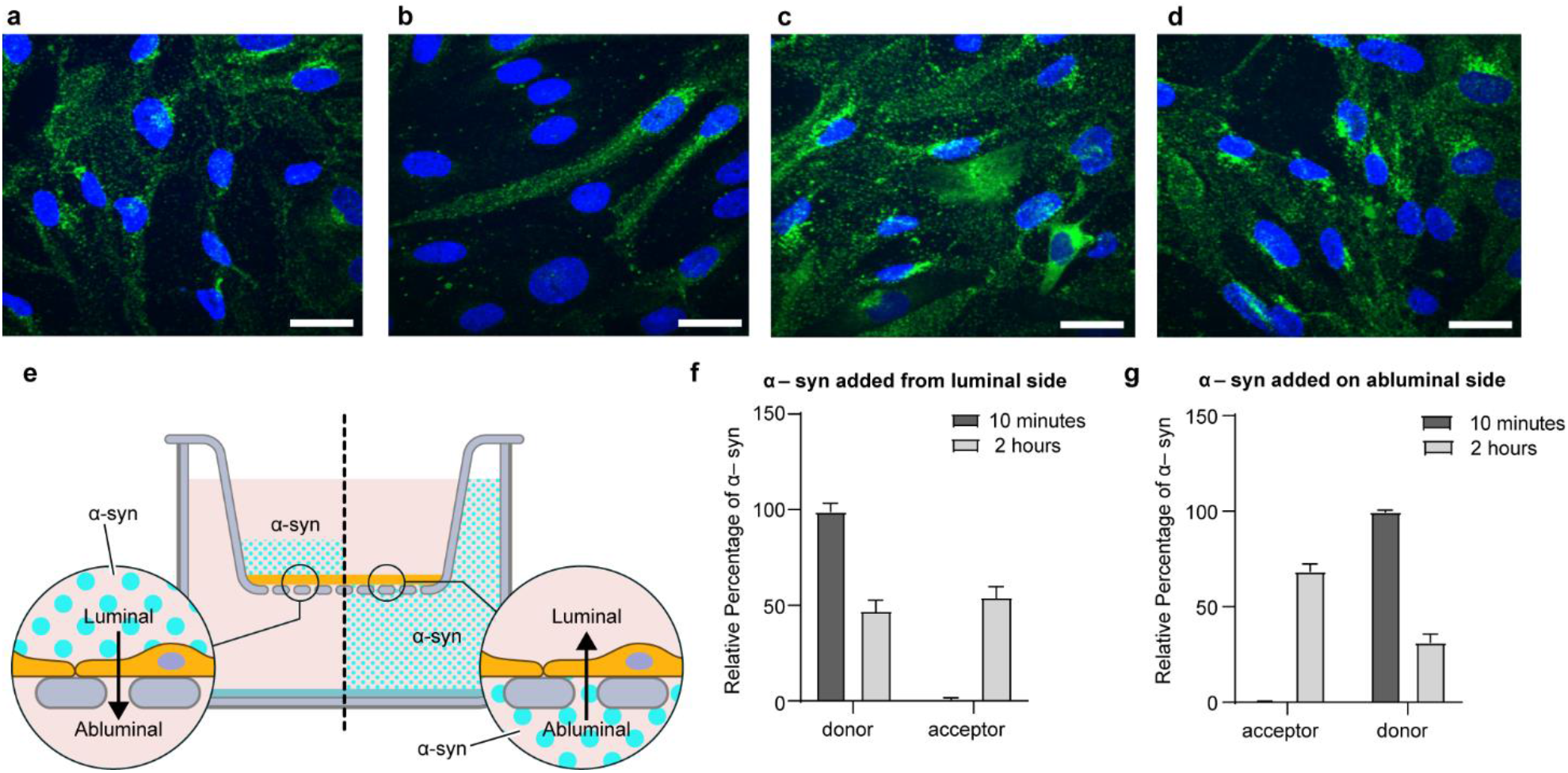
Transport of α-syn monomers through the BBB model. Representative confocal micrographs of α-syn monomer added to pBECs seeded on filters. Green are stains for the α-syn monomer and blue are Hoechst stains to mark the nucleus of the cells. (a) 10 min luminal side (b) 10 min abluminal side (c) 2 h luminal side and (d) 2 h abluminal side, scale bars show 15 µm. (e) Experimental set up: 100 nM α-syn monomer was added on to the BBB cell model and incubated for 10 min or 2 h at 37 °C with shaking at 150 rpm (f) ELISA results for α-syn transport across BBB when added from the luminal side (g) ELISA results for α-syn monomeric transport across BBB when added from the abluminal side.

Although the difference is relatively modest, it suggests different mechanisms of α-syn transport in pBECs when trafficking from luminal or abluminal site of pBECs. In addition, we also studied the uptake of α-syn oligomers to investigate whether these are capable of crossing the BBB model. Imaging showed that α-syn oligomers were endocytosed by pBECs from either side of the BBB model, though with a higher degree of endocytosis or intracellular accumulation when added from abluminal side (**Fig. 3a-b**). In contrast to monomers, only a small amount of α-syn oligomers was measured to transcytose through the cell layer after 2 h (**Fig. 3c**). The fraction succeeding transcytosis occurred in the luminal to abluminal direction (**Fig. 3c**), suggesting accumulation or stalled trafficking of the oligomeric form of α-syn in the abluminal to luminal direction. Due to the limited trafficking of the oligomeric form through our BBB model, we chose to conduct all the subsequent trafficking experiments with α-syn monomers.

**Figure 3:**
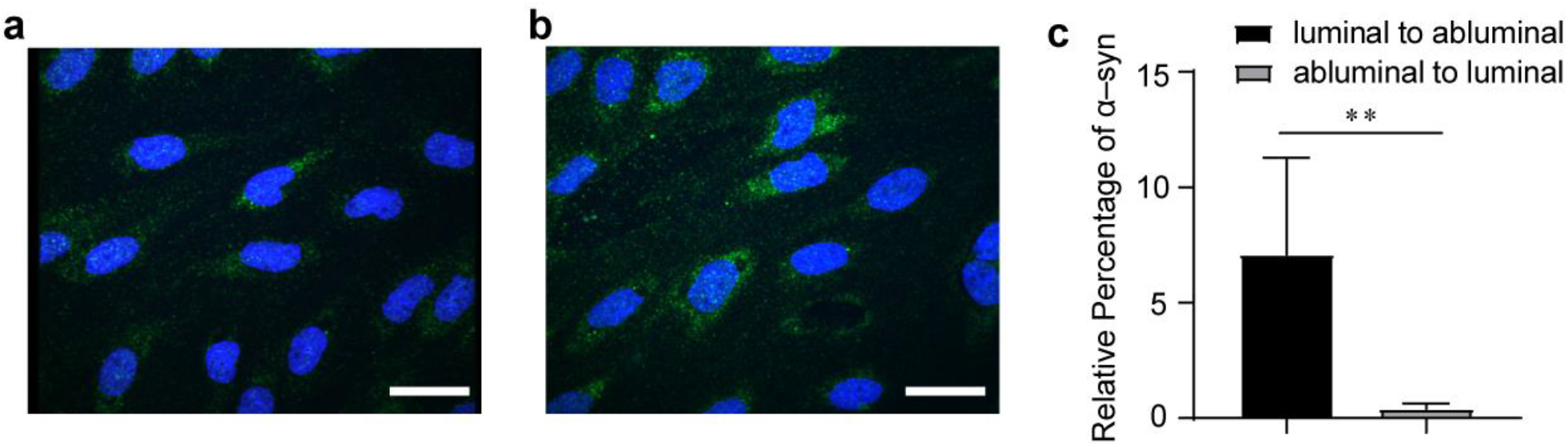
Transport of α-syn oligomers through the BBB model. Representative confocal microscopy images of α-syn ONE oligomers uptake in pBEC on filters after 2 h chase. Green color are stains for α-syn oligomers and blue are Hoechst stains. (a) α-syn oligomer added from luminal side (b) α-syn oligomer added from abluminal side (c). Scale bars show 15 µm. ELISA results for α-syn oligomeric transport across BBB with 2 h of chase for the indicated directions. The statistical test used was Welch’s t test (**P < 0.01) error bars are Standard deviations.

### Intracellular trafficking route of α-syn monomers in pBECs

To ensure that the purified monomeric α-syn did not affect the integrity of the pBEC barrier, we assessed the effect of α-syn on permeability using the similar sized 14 kD FITC dextran as tracer. To this end, pBECs were incubated with α-syn (+/-) and 14 kD FITC dextran for 2 h. As shown in **Supplementary Table S3**, there was no significant difference in permeability of FITC dextran between α-syn treated and untreated pBECs. This rules out paracellular trafficking and favors intracellular trafficking of α-syn through the applied BBB model.

In order to establish the intracellular trafficking route of α-syn in endothelial cells in NCC astrocyte co-culture, we performed colocalization studies between monomeric α-syn and a selection of endothelial trafficking machinery (33). We chose to stain for trafficking along a putative endocytic-exocytic cellular trafficking path entailing clathrin (marker for clathrin mediated endocytosis and vesicular sorting), early endosomal antigen1 (EEA1), VPS35 (marker for retrograde sorting), Rab7 (marker for late endosome/lysozyme sorting) and Rab8a (Golgi-exocytic sorting). We also included caveolin1 in the analysis because this protein has been proposed to be involved in regulating endothelial transcytosis (34, 35). Immunofluorescent colocalization images for α-syn and EEA1 from the luminal side after 10 min of α-syn treatment are shown in **Fig. 4**. Immunofluorescence colocalization images for all other markers are shown in **Supplementary Fig. S2**. Colocalization data from for all six markers are summarized in **Fig. 4b**, showing very similar colocalization between α-syn and each marker after 10 min and 2 h of α-syn chase, irrespective whether α-syn was added to the luminal or abluminal side of the BBB. However, the level of colocalization varied among the different markers. There was strong colocalization of α-syn with clathrin after 10 min of chase both from the luminal and abluminal side of the BBB model, indicating that α-syn is internalized in a clathrin-dependent manner in BECs. We observed weak colocalization of α-syn with caveolin1, suggesting α-syn internalization independent of caveolin1 in pBECs. Some colocalization occurred between α-syn and EEA1, and was strong between α-syn, Rab7 and VSP35. Rab8a did not colocalize with α-syn at any time. Collectively, these results suggest that α-syn is internalized in a clathrin-dependent manner and preferentially sorts through the cell via a late endosome retromer-connected trafficking pathway.

**Figure 4:**
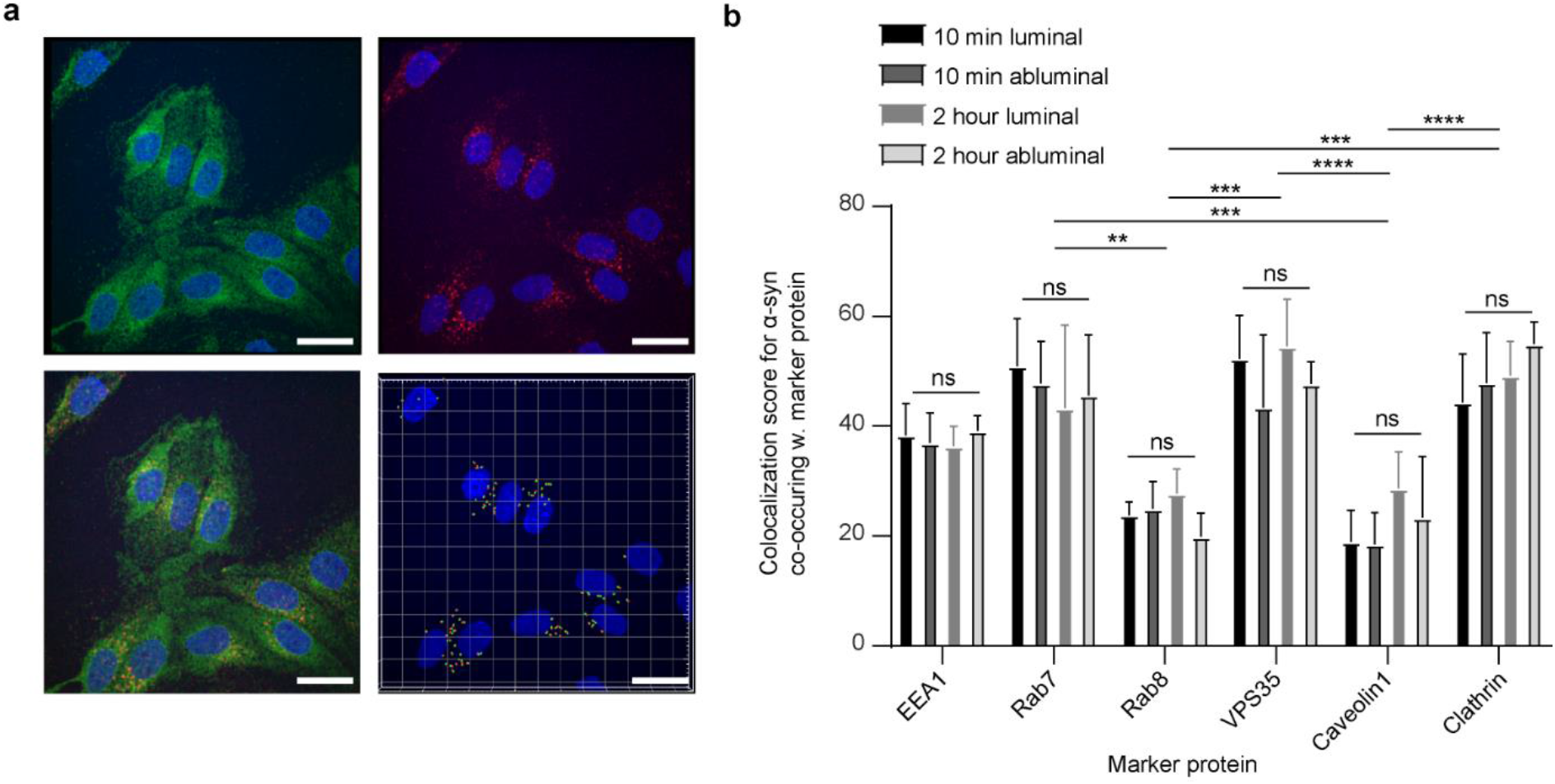
Analysis of α-syn co-occurrence with trafficking markers. Representative confocal micrographs of α-syn monomer treated pBECs on filters with α -syn added for 10 min to the luminal side (a). Green show α-syn monomer stain, blue is Hoechst stain and red is the EEA1 stain. Lower right micrograph shows the outcome of IMARIS spot segmentation and colocalization analysis between α-syn and EEA1 channels. Scale bars show 15 µm. (b) Semi-quantification of α-syn co-occurrence with different compartments in pBEC: EEA1 (early endosome), RAB7 (endosome-lysosome), VPS35 (endosome-golgi, retromer), Rab8a (Golgi-plasma membrane), Caveolin1 (Caveolae/transendothelial channels) and Clathrin (endo-lysosomal vesicles). The statistical test used was Tukey’s multiple comparisons test (**P < 0.05, *** P< 0.001 and **** P< 0.0001) error bars are Standard deviations

### Rab7 and VPS35 regulate monomeric α-syn intracellular transport in pBECs

To further validate the colocalization analysis showing Rab7/VPS35 directed trafficking of monomeric α-syn across the BBB model, we knocked down Rab7 and VPS35 using small interfering RNAs (siRNA) and analyzed the effect on α-syn trafficking. We confirmed protein knock down by western blotting (**Fig. 5d**) and analyzed uptake and transport of α-syn from 2 h α-syn chase treatment.

**Figure 5:**
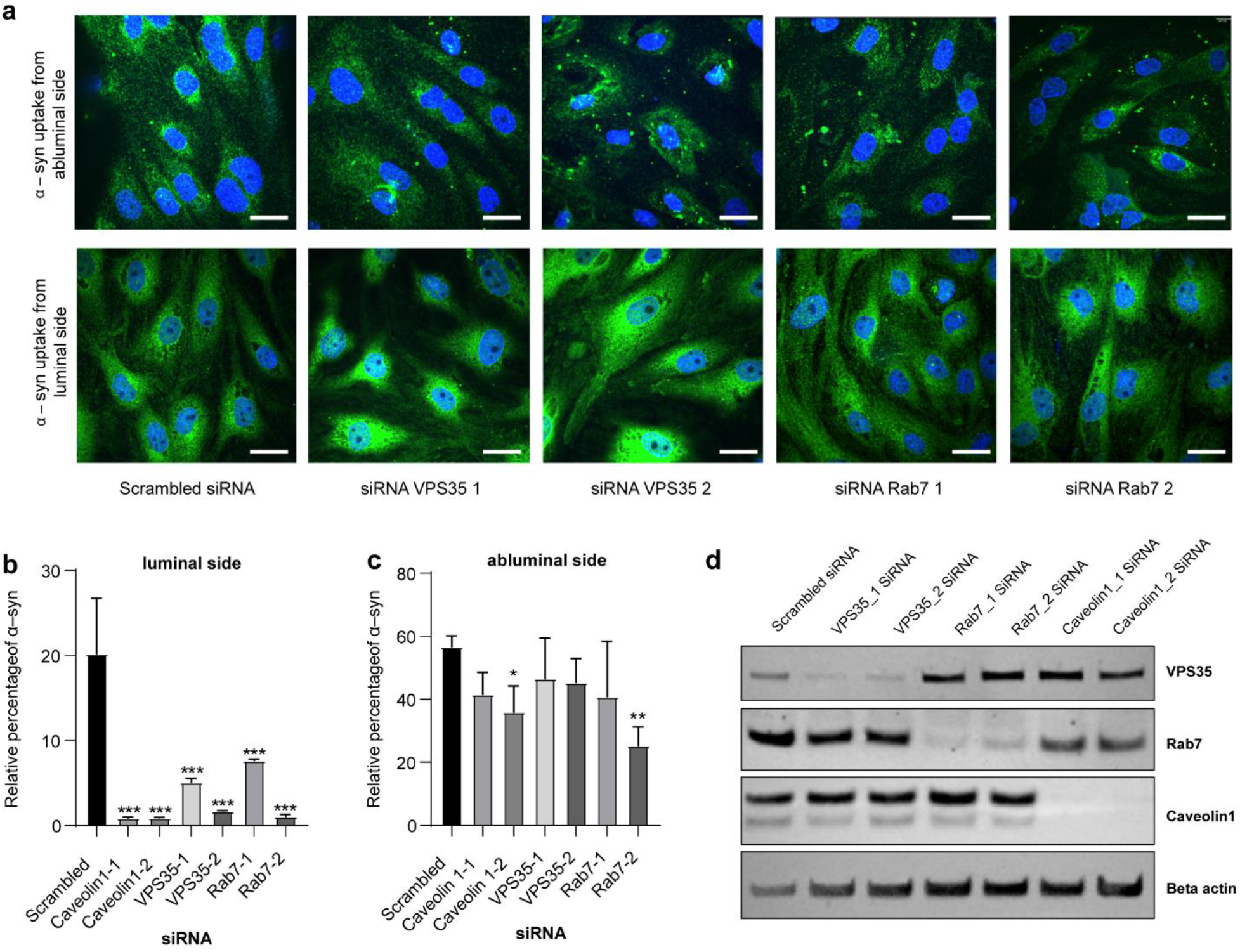
Effect of intracellular trafficking machinery on α-syn transport through the BBB model. Representative micrographs of α-syn uptake in pBEC’s on filters knocked down for VPS35 and Rab7 and treated for 2 h with monomeric α -syn, as indicated in (a). Scale bars show 15 µm. Relative percentage of α-syn transport in cells pretreated with indicated siRNA (in comparison to scrambled siRNA) and treated for 2 h with α-syn from luminal (b) and abluminal side of the BBB model (c), respectively, The statistical test used was a Dunnett’s multiple comparisons test (*P < 0.05, ** P< 0.01 and ** P< 0.001) error bars are Standard deviations. Representative Western blot to confirm the efficiency of the siRNA mediated knockdown of indicated proteins in pBECs on Transwell filters.

We found that α-syn is taken up by pBECs regardless of Rab7 and VPS35 knockdown (**Fig. 5a**). Uptake occurred regardless of whether α-syn was added to pBECs from the luminal or abluminal side. However, ELISA showed that knockdown of Rab7/VPS35 almost abolished transport of α-syn from the luminal to the abluminal side (**Fig. 5b**). In contrast, there was significant transport of α-syn from the abluminal to the luminal side after Rab7/VPS35 knockdown (**Fig. 5c**).

Caveolin1 has been found to be involved in regulating transcytosis of different components in BBB, suggesting a global effect of caveolin1 on BEC transcytosis (34, 35). We similarly tested if Caveolin1 KD could affect α-syn transcytosis and found that Caveolin1 KD also abolished luminal to abluminal α-syn transcytosis (**Fig. 5b**). Interestingly, our colocalization data showed that α-syn did not colocalize with caveolin1 (**Fig. 4** and **Fig. S2**), suggesting an indirect role of caveolin1 on α-syn trafficking.

While colocalization and ELISA data showed an indirect role of caveolin1 in α-syn transcytosis, our colocalization and ELISA data indicate direct roles for both Rab7 and VPS35 α-syn trafficking (**Fig. 4-5**). To test this, we analyzed the colocalization of α-syn with Rab7 and VPS35 in greater detail using STED microscopy. The STED Imaging showed that α-syn was positioned in carriers decorated with Rab7 but containing no VPS35 (**Fig. 6a-b and 6d-e**). Quantification of the Pearson correlation coefficients (PC) obtained from STED images of co-stains for α-syn confirmed a high level of colocalization between α-syn and Rab7 (PC ∼ 0.5) but almost no colocalization with VPS35 (PC ∼0.2) as shown in **Fig. 6c** and **6f**. Overall, these results show that Rab7 might perform a direct role in luminal to abluminal α-syn trafficking whereas VPS35 function is secondary.

**Figure 6:**
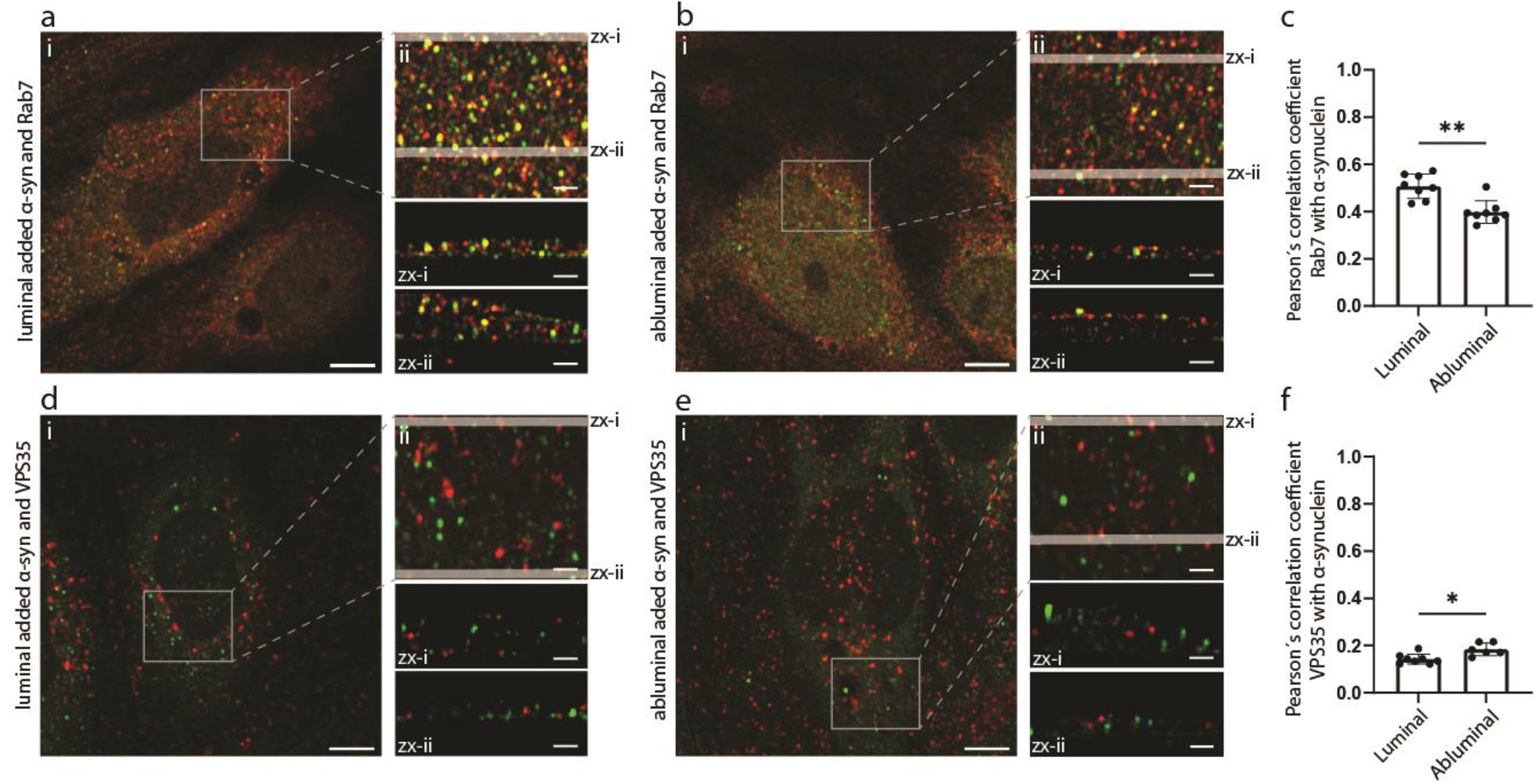
Rab7 decorates α-syn carriers. Quantification of Rab7 and VPS35 co-localization with α-syn added from the luminal or abluminal side. Panels (a) and (b) are micrographs of pBEC cells with luminally and abluminally added α-syn, respectively, co-stained for Rab7. Panels d and e are micrographs of pBEC cells with luminally and abluminally added α-syn, respectively, co-stained for VPS35. Representative dual-color confocal micrograph (i) with Rab7 or VPS35 (red) and α-syn (green), scale bar show 5 µm. Maximum z-projected 3D STED stack (ii) of the highlighted area in (i), scale bar show 1 µm. Micrographs (zx-i) and (zx-ii) are sideview representations of the cells from maximum y-projections of the 400 nm slices highlighted in (ii). Scale bar show 1 µm. All images are contrast adjusted to enhance clarity. Pearson’s correlation coefficients between Rab7 and α-syn are shown in (c) and between VPS35 and α-syn in (f).The statistical test used was an unpaired Mann-Whitney test (\**p* < 0.05 and ** *p*< 0.01). Error bars are standard deviations.

## Discussion

Understanding the transport of α-syn monomers and oligomers from blood to brain or vice versa across the BBB is essential to understand the potential role of peripheral α-syn species in PD pathology. Currently, emerging questions are if aggregates and PD propagation could be stopped by targeting peripheral α-syn or perhaps a component of the transport machinery in the endothelial cell layer of the BBB?

While the apicobasal cell polarity causing polarized transport in brain endothelial cells is well known (36), the machinery involved in regulating this trafficking is largely unknown. Therefore, it is highly warranted to establish more knowledge and here we report the usability of our BBB model setup to study the trafficking machinery behind such polarized transport of α-syn.

In this study, we added purified α-syn to the cellular medium of an *in vitro* Transwell BBB model to mimic extracellular uptake from blood stream to brain parenchymal fluid, or vice versa. Our experimental setup shows both binding and uptake from luminal and abluminal cell surfaces (**Fig. 2a-c**). In addition, uptake from either side leads to transcytosis of extracellular α-syn through the pBEC cell layer (**Fig. 2f-g**). Such bidirectional transport was also reported in *in vivo* models (23) and has been observed for other molecules such as potassium and glucose, whose CNS levels are tightly regulated in order to maintain brain homeostasis.

The oligomeric form of α-syn has a larger surface for binding to cellular surfaces when extracellular transport occurs. This makes it reasonable to expect that other cellular trafficking paths are involved for its transport than for the monomeric form. When we added α-syn oligomers to the medium, binding and uptake was low but evident from both sides, though with more oligomeric α-syn apparent when added from the abluminal side (**Fig. 3a-b**). Transport of oligomeric α-syn through the pBEC layer was however only measurable from the luminal to abluminal side of our BBB model (**Fig. 3c**), suggesting selective transport from luminal to abluminal side and intracellular accumulation when added from the abluminal side (**Fig. 3b-c**). Polarized transport is a common biological mechanism in BECs (36) and here we show that it causes a trap for oligomeric α-syn when we mimic brain to blood transport using our BBB model. This observation is important in order to understand the accumulation of oligomeric α-syn in brain. Reports suggest that there is an association between α-syn aggregation and endothelial cell degeneration in PD pathology (37, 38). Currently there are only few *in vitro* BBB models (39) to study the biology behind this mechanism and it is unknown if these models can provide the correct setup to mimic the suggested polarization trap.

We observed no polarized trafficking of monomeric α-syn; rather it followed a clathrin - early endosome - retromer - late endosome directed trafficking path. The path allowed transcytosis of α-syn through the tight pBEC layer, but there was no indication of exit via Golgi-plasma membrane transport, indicated by the Rab8a colocalization data (**Fig. 4)**.

Both Rab7 and VPS35 are associated with accumulation of α-syn and VPS35 is associated with PD pathology (40, 41). Dinter *et al* showed the protective role of Rab7 overexpression in HEK293 cells and in Drosophila model (40). HEK293 cells expressing A53T α-syn, overexpression of Rab7 reduced the percentage of cells with α-syn particles and the overall amount of α-syn in cells. Overexpression of Rab7 protects locomotor deficit induced by neuronal expression of A53T α-syn in Drosophila. This protective role of Rab7 could be attributed to its role in autophagy (42). Knockdown of *VPS35* in *Drosophila expressing human* α-syn induced the accumulation of α-syn species in the brain and exacerbated both locomotor impairments and mild compound eye disorganization (43). Overexpression of VPS35 has been reported to ameliorate the pathogenic mutant LRRK2 eye phenotype in Drosophila PD model (44). Recently, Chen *et al* developed D620N VPS35 knock in mice model for inherited PD that develops robust and progressive degeneration of nigral dopaminergic neurons (45). Furthermore, recent reports show that an increase in Caveolin1 expression in aged individuals and in aged mice correlates with increased uptake of α-syn (46, 47). We therefore used siRNA to remove these proteins from the BBB cell model to study their effect on α-syn trafficking. Strikingly, all of the proteins affected α-syn transcytosis from the luminal side to the abluminal side but not the opposite direction (**Fig. 5**). This suggests that there are two different pathways involved in facilitating the α-syn transport, *i*.*e*. an unknown path for abluminal to luminal transport and a path dependent on Rab7, VPS35 and caveolin1 expression for luminal to abluminal transport. We further used STED microscopy to dissect the protein associations and found that only Rab7 marked the α-syn containing membrane carriers. Interestingly, a recent study also found transcytotic carriers decorated with Rab7 (48), suggesting it as a possible marker of luminal to abluminal transcytotic carriers. The activity of Rab7 is known to be regulated by the VPS35 retromer complex (49), which explains VPS35s indirect role in regulating α-syn transcytosis in our BBB model. We were unable to explain why Caveolin1 affects luminal to abluminal α-syn transcytosis. Since the protein does not colocalize with α-syn (**Fig. 4** and **Fig. S2**), it must perform an indirect function in regulating the polarized trafficking path. A study in mouse fibroblasts suggest a role of caveolin1 in regulating cell polarity (50), which could be a plausible explanation for its effect on the polarized trafficking we find in our pBEC.

In conclusion, we report a functional BBB model, which show bidirectional transport of α-syn monomers and polarized transport of α-syn oligomers. We propose that this model can be used to study the mechanism of accumulation and aggregation of α-syn in BECs, which currently remains an enigma in PD pathology. Using this model, we find that Rab7 regulate transcytic carriers essential for transporting monomeric α-syn through from luminal to abluminal side of pBECs. Furthermore, our experiments identify the presence of one or two unknown polarized trafficking paths directing trafficking of monomeric and oligomeric α-syn from abluminal to luminal side of pBECs. In this direction, monomeric α-syn transcytoses through the cells, while the oligomeric form is accumulating inside the cells. Understanding the regulation of these carriers can be expected to be important for future alleviation of brain accumulation of α-syn.

## Materials and Methods

### Protein expression and purification

α-syn was expressed in *E. coli* and purified as described previously (29). For all experiments, fresh samples were prepared by dissolving lyophilized α-syn in phosphate saline buffer (PBS) (20 mM phosphate, 150 mM NaCl, pH 7.4) and filtered (0.2 μm) prior to use. The concentration was determined with a Nano Drop (ND-1000, Thermo Scientific, USA) using a theoretical extinction coefficient of 0.412 (mg/mL) ^−1^ cm^−1^.

### Purification of pBECs

Porcine brain capillaries were purified based on established protocols (28). In brief, grey matter from six-month old animals was isolated and homogenized using a 40 mL Dounce tissue grinder (DWK Life Sciences GmbH). The homogenate was filtered through 140 μm mesh filter (Cat no.: NY4H04700, Merck Millipore, KGaA, Darmstadt, Germany). The filtrate was enzymatically digested for 1 h with 500 μg ml-1 Collagenase type II (Cat no.: 17101-015, Gibco, Grand Island, NY)), 2.5% Trypsin-EDTA (1:10 dilution;,Cat no.: Thermo Fisher Scientific, 2665 NN BLEISWIJK NETHERLANDS 15090-046) and 50 μg ml-1 DNAse I (Cat no.: D4513, Sigma-Aldrich, GmbH Mannheim Germany). Digested capillaries were collected by centrifugation at 240 rcf, 4 °C for 5 min.

### Establishment of BBB model

Following isolation, porcine brain capillaries were seeded in T75 flasks (Thermo Fisher Scientific, Roskilde, Denmark) coated with collagen IV (500 μg/ml) and fibronectin (100 μg/ml) in DMEM/F12 medium supplemented with 10% plasma-derived bovine serum (PDS; First Link Ltd., Wolverhampton, UK), PS (Penicillin (100 U/mL) streptomycin (100 µg/mL)), (15 U/mL) heparin and puromycin. Cultivation of pBECs with (4 µg/mL) puromycin was continued for four days before reseeding pBECs without puromycin on Transwell filters. Isolation of primary rat astrocytes was carried out as previously described [25], and afterwards cultured in a 12-well plate coated with poly-L-lysine (5 μg/ml) in ddH_2_O. 24 h before seeding pBECs on inserts, astrocyte cultures were refreshed with 1.5 mL medium, allowing time for astrocytic release of factors inducing the endothelial phenotype and barrier integrity. The non-contact co-culture (NCC) was established by seeding 1.1×10^5^ pBECs in 500 µL of pBEC media per Transwell filter (Cat no: 3401, Corning, Kennebunk ME 04043 USA), which were afterwards transferred to wells with pre-cultured primary rat astrocytes. 3 days post NCC establishment, endothelial- and astrocytic cells were further stimulated with the differentiation factors cAMP (250 µM), hydrocortisone (550 nM), and RO (17 µM) to increase the barrier development. The barrier development was evaluated by measuring transendothelial electrical resistance (TEER), with values above 1000 Ω*cm2 applied for experiments. During experiments, TEER values were measured every 24 h using EndOhm-12 and EVOM measurement device (World Precision Instruments).

### Treatment of pBECs with α-synuclein

After measuring the TEER values, Transwell inserts with pBECs were transferred to new plates without astrocytes, and containing only serum free medium. Medium within Transwell inserts was replaced with fresh medium and incubated for 30 min in incubator. 100 nM α-syn was then added to the luminal or abluminal compartment of the Transwell system, and kept at 37 °C in the incubator while shaking at 150 rpm. Medium was collected at two time points (10 min and 2 h), freeze stored at −20 °C for later ELISA measurements and Transwell filters were fixed for immunostaining.

### Enzyme linked immunosorbent assay (ELISA) assay

96-well Maxisorp plates were incubated with 100 μl 2 µg/ml ASY-1 in ELISA coating buffer (0.1MNaHCO_3_ in phosphate buffered saline PBS (2.8 mM NaH_2_PO_4_, 7.2 mM Na_2_HPO_4_, 123 mM NaCl), pH = 9.5) overnight (30). The plate was washed three times with TBS-Tween (1.5M NaCl, 200 mM Tris, 0.05% Tween-20, pH 7.4) and incubated with ELISA blocking buffer (PBS + 10% fetal calf serum (FCS)) for 2 h at room temperature (RT). After washing, 100 μl medium samples collected from pBECs after 10 min and two hrs (see previous sections) were added to ELISA wells and incubated overnight at 4 °C. The following day, wells were washed, and 100 μl 0.5 μg/ml mouse monoclonal anti-α-syn (BD Transduction Laboratories, 610787) was added for 2 h. After washing, anti-mouse-HRP antibody was added for 30 min followed by wash and addition of 100 μl 3,3’,5,5’-Tetramethylbenzidine to each well for 15 min. Finally, the enzymatic reaction was stopped by the addition of 100 μl 1M phosphoric acid. The absorbance was measured at 450 nm and compared to signals for α-syn standard curves.

### Small interfering RNA (siRNA) transfection

pBECs were generated as described above, including four days of puromycin to obtain endothelial-selection (28). On the fourth day, pBECs were re-seeded on collagen IV and fibronectin coated 6-well plates with a density of 8.2 × 10^4^ cells per well in complete medium containing DMEM/SF9 media, supplemented with 10% plasma derived serum (PDS), 50 U/ml penicillin and 50µg/ml streptomycin and 15U/ml heparin. On day five, medium was exchanged to DMEM with 10% PDS and cells were transfected with 40 nM siRNA using lipofectamine 3000 (Invitrogen). On day six, medium was renewed and a consecutive siRNA transfection was done 5 h before seeding cells on Transwell filters. The siRNAs were purchased from Integrated DNA Technologies (IDT, GmbH) targeting VPS35 sequences: CCTGACAGATGAGTTTGCTAAAGGA and GGATTCGCTTCACACTGCCACCTTT, RAB7a sequences: AATCAGATCTTTTTACAGTAUCCAT and CTGGTGCTACAGCAAAAACAACATT, and Caveolin1 sequences: GAA TGAGGTCAGCATGTCTATTCAG and CATCAGCCGTGTCTATTCCATCTAC. IDT’s scrambled negative control DsiRNA was used for control transfection.

### Antibodies

All antibodies are commercially available (except ASY-1, which was produced in-house as a polyclonal antibody (30). A list of primary and secondary antibodies are given in Supplementary Table 1 and 2.

### Immunostaining and Confocal Microscopy

Samples were fixed with cytoskeleton fixation buffer containing 10 mM MES, 3 mM MgCl_2_, 138 mM KCl, 2 mM EGTA, 0.32 M sucrose and 4 % PFA for 20 min at room temperature. Permeabilization and blocking were done with 0.1% Triton-X100 and 2% bovine serum albumin in PBS. Primary antibodies were diluted 1:200 in blocking solution and incubated with the samples for 1h at RT. Samples were treated with secondary antibodies at 1:500 dilutions for 30 min at RT. For staining of the nuclei, samples were incubated with Hoechst 32528 in distilled water (0.5 μg/ml). The samples were mounted on glass slides using Dako fluorescence mounting medium (Dako, Glostrup, Denmark). Confocal images were captured by Olympus IX-83 fluorescent microscope with a Yokogawa CSU-X1 confocal spinning unit and Andor iXon Ultra 897 camera, Olympus UPLSAPO W, ×60/ 1.20 NA water objective lens, using Olympus CellSens software (Olympus). Images were processed using Imaris software. For each channel, laser power was adjusted and applied independently.

### STED microscopy

Dual-color 3D stimulated emission depletion (STED) image acquisition was carried out on an Abberior Facility Line STED microscope using a 100x magnification UplanSApo 1.4 NA oil immersion objective lens. Sequential imaging of Abberior STAR ORANGE (STAR ORANGE) and Abberior STAR RED (STAR RED) using a pulsed excitation laser of either 561 nm or 640 nm, respectively, and for both dyes a pulsed 775 nm depletion laser was used. Fluorescence was detected by spectral detectors between 580-630 nm or 680-763 nm with a time gating delay of 750 ps for an interval of 8 ns. Image stacks were recorded with a pixel size of either 60 or 40 nm in x, y, and z.

### Image analysis

Colocalization score was analyzed and measured as described previously using IMARIS software (Bitplane) for spot segmentation (31). Image analysis of the 3D STED microscopy images was carried out in MATLAB 2020b. Each image stack was smoothened with a 3D median filter of 3×3×3 pixels to minimize background noise. Due to inhomogeneous auto fluorescence from the polycarbonate filter membrane on which the cells were cultured, we performed a local background correction on each xz-slice of the image stack by two morphological operations, an erosion and a dilation, with a disk size of five pixels. To estimate the amount of colocalization between α-syn and either Rab7 or VPS35, we determined the Pearson correlation coefficient (32). To minimize background contribution, each cell was divided into small sections of 400 nm along y, and maximum intensity projections were calculated along y for the intervals. For each slice, the Pearson correlation coefficient was calculated by the following equation:

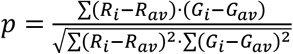

Where *R*_*i*_ is the pixel intensity in each pixel and *R*_*av*_ is the mean intensity of the slice for the STAR RED signal. *G*_*i*_ and *G*_*av*_ are corresponding intensities for the STAR ORANGE signal. To obtain the Pearson correlation coefficient for each cell, the mean for the slices was calculated.

## Supporting information

supplementary information Alam et al 2021

## Acknowledgments

We are thankful to the Novo Nordisk Foundation for supporting P.A., M.S.N and D.E.O. (grant NNF17OC0028806). We are grateful to Annemette Boe Marnow for her skilled technical support and Anita Mora, graphics art unit at RML for graphics assistance.

## Notes

### Competing Interest Statement

The authors have declared no competing interest.

