## supplementary information Alam et al 2021 for "Polarized α-synuclein trafficking and transcytosis across Brain Endothelial Cells via Rab7-decorated carriers"

[†]Equal contributors

**Supplementary Table 1: List of primary antibodies**

| Target protein | Antibody | Manufacturer | Cat. No. |
|---|---|---|---|
| α- synuclein | Rabbit polyclonal anti α-syn (ASY-1) | Poul Henning Jensen laboratory | |
| α- synuclein | Mouse monoclonal anti α-syn | | 610787 |
| EEA1 | Mouse polyclonal anti EEA1 | BD biosciences | 610547 |
| Caveolin 1 | Rabbit polyclonal anti cav1 | st. John's lab | STJ92051 |
| Rab 7 | Mouse monoclonal anti Rab7 | Abcam | ab50533 |
| VPS35 | Goat polyclonal anti VPS35 | Everest Biotech | EB06268 |
| Clathrin | Mouse anti clathrin | Lundbeck, Denmark | |
| Rab 8 | Rabbit monoclonal anti Rab8 | Cell signaling | CST-6975T |
| Claudin 5 | Mouse monoclonal anti claudin 5 | Thermo Fisher | 35-2500 |
| ZO1 | Rabbit polyclonal anti ZO1 | InvitrogenInvitrogen | 61-7300 |

**Supplementary Table 2: List of secondary antibodies**

| **Applied for** | **Antibody** | **Manufacturer** | **Cat. No.** |
|---|---|---|---|
| Immunofluorescence labelling of α-syn | Donkey-Anti mouse Alexa 488 | Invitrogen | A21202 |
| Immunofluorescence labelling of VPS35 | Donkey-anti Goat Alexa 647 | Molecular probes | A21082 |
| Immunofluorescence labelling of Rab8a, caveolin 1 | Goat-anti rabbit Alexa 647 | Invitrogen (Molecular probes) | A21244 |
| Immunofluorescence labelling of α-syn | Donkey-Anti rabbit Alexa 488 | Invitrogen (Molecular probes) | A21206 |
| Immunofluorescence labelling of EEA, Rab7, clathrin | Goat-anti mouse Alexa 647 | Invitrogen (Molecular probes) | A21235 |
| Immunofluorescence labelling of α-syn | Donkey Anti Rabbit STRAR ORANGE | Invitrogen | A16031/1 mg Abberior Star Orange-NHS ester |
| Immunofluorescence labelling of Rab7 | Goat Anti Mouse STAR RED | Abberior, GmbH | STRED-1001-500UG |
| Immunofluorescence labelling of VPS35 | D Anti G STAR RED | Abberior. GmbH | TRED-1055-500UG |

**Supplementary table 3:** Apparent permeability coefficient (P app cm/s) of 14 kD FITC- dextran across BBB

| | **Average Papp*x10^-6^*** | **SD** |
|---|---|---|
| Dextran + α- syn | 2.21 | 0.26 |
| Dextran | 6.03 | 0.56 |

**Supplementary figures**

**Fig. S1**

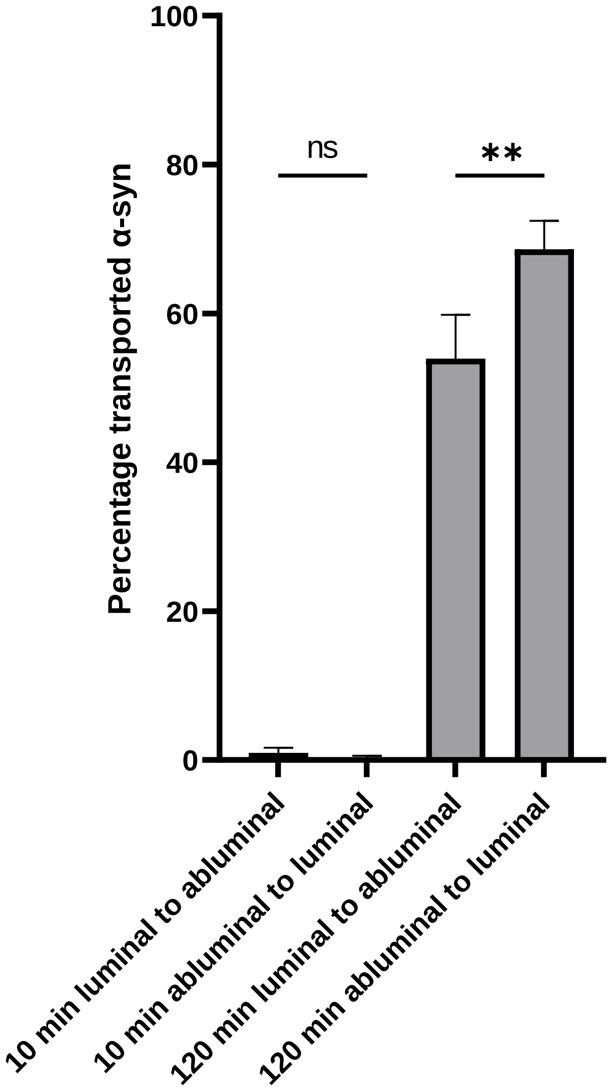


**Supplementary figure 1: The amount of transported monomeric α-syn is significantly different depending on transport direction.** In relation to main figure 2, statistical difference was measured using ordinary one-way ANOVA (**$P < 0.01$) error bars are Standard deviations.

**Fig. S2**

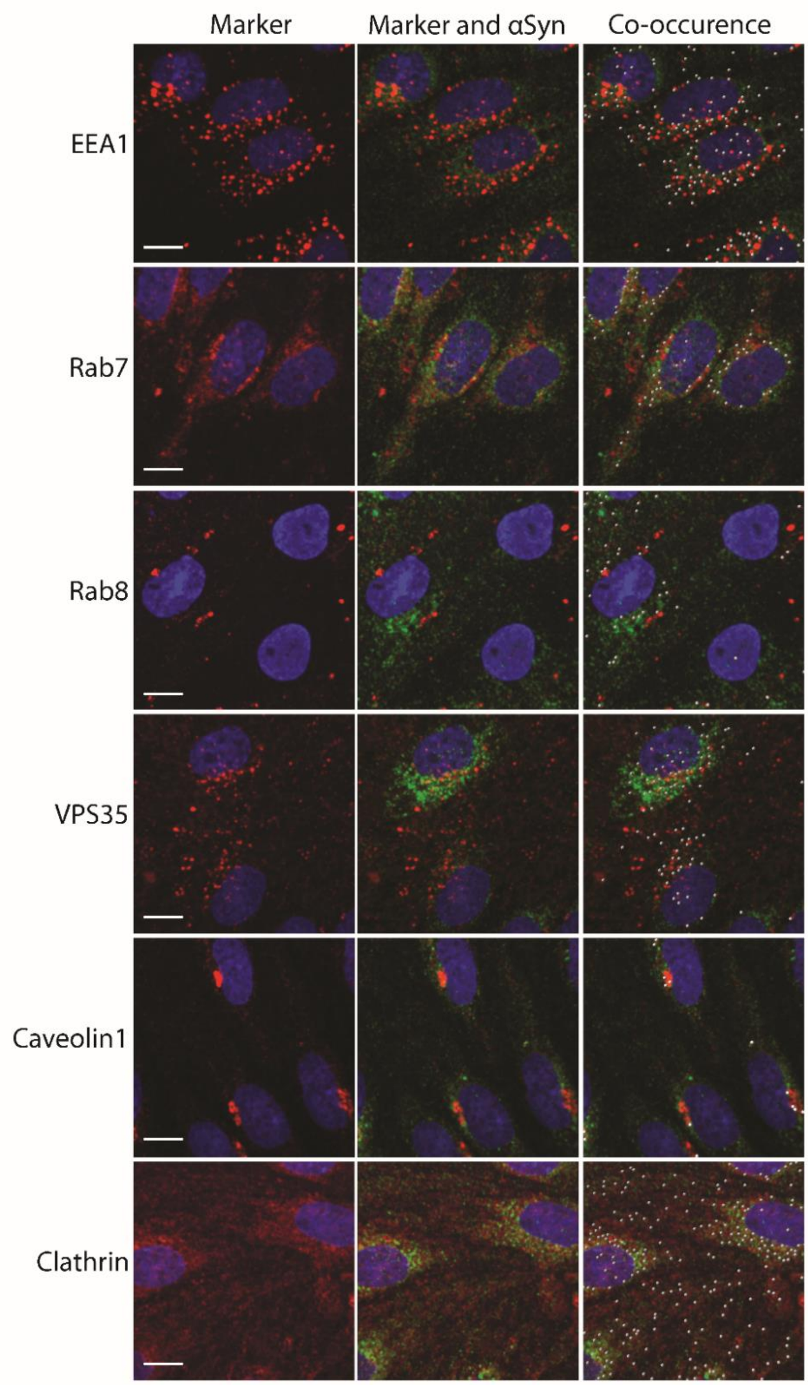


**Supplementary figure 2: Representative stains for α-syn co-occurrence with trafficking markers.** Representative Maximum projected 3D stacks from confocal micrographs of α-syn monomer treated pBECs on filters with α-syn added for two hours to the luminal side. Green show α-syn monomer stain, blue is Hoechst stain and red is the indicated marker stain. Right micrograph in panels shows segmented IMARIS spots of colocalization analysis between α-syn and marker channels with with white points indicating co-occurence. Scale bars show 10 μm.
